# Tracer-based metabolomics for profiling nitric oxide metabolites in a 3D microvessel-on-a-chip model

**DOI:** 10.1101/2023.12.03.569402

**Authors:** Kanchana Pandian, Luojiao Huang, Abidemi Junaid, Amy Harms, Anton Jan van Zonneveld, Thomas Hankemeier

## Abstract

Endothelial dysfunction is a common denominator in cardiovascular diseases (CVDs) associated with diabetes, hypertension, obesity, renal failure or hypercholesterolemia. In these disease states, circulating adverse metabolic or hemostatic risk factors drive the progression of inflammation, thrombosis, platelet activation and atherosclerosis. A hallmark of endothelial dysfunction is the reduced bioavailability of nitric oxide (NO), a signaling molecule essential for vascular homeostasis. Numerous studies have focused on NO synthesis by endothelial cells (ECs) using *in vitro* cultures to understand the pathophysiology of endothelial dysfunction. A limitation of these studies is that the expression of the NO-generating enzyme, endothelial nitric oxide synthase (eNOS), in physiological conditions is modulated by the exposure of the ECs to laminar shear stress, a stimulus that is clearly lacking in most two-dimensional (2D) cultures.

Here we developed a tracer-based metabolomics approach to measure NO-specific metabolites with mass spectrometry (MS) and show the impact of unidirectional fluid flow on metabolic parameters associated with NO synthesis using 2D and three-dimensional (3D) platforms. Specifically, we tracked the conversion of stable-isotope labeled NO substrate L-Arginine to L-Citrulline and L-Ornithine to determine eNOS activity. We demonstrated that when human coronary artery endothelial cells (HCAECs) cultured in media containing ^13^C_6_,^15^N_4_-L-Arginine treated with eNOS stimulator – vascular endothelial growth factor (VEGF), eNOS inhibitor – L-NAME and arginase inhibitor - S-(2- boronoethyl)-L-cysteine (BEC), their downstream metabolites - ^13^C_6_,^15^N_3_ L-Citrulline and ^13^C_5_,^15^N_2_ L-Ornithine showed clear responses as measured using Ultra-performance liquid chromatography tandem mass spectrometry (UPLC-MS/MS). In this study, we also assessed the NO metabolic status of a static 2D culture, a 3D microvessel model with bidirectional flow, and our 3D model with unidirectional fluid flow generated by a microfluidic pump. Compared to 2D culture, our 3D model showed significant effects in the control and microvessels exposed to VEGF when Citrulline/Ornithine ratio was analyzed. The obtained result indicates that the 2D static culture mimics more endothelial dysfunction status. Our detection method and 3D model with a unidirectional fluid flow provides a more representative physiological environment that exhibits perfect model to study endothelial dysfunction.

## Introduction

Nitric Oxide is an essential diatomic molecule that is generated by endothelial nitric oxide synthase (eNOS/NOS3) and that exerts multiple key roles in vascular homeostasis, including vasodilation and inflammation. The eNOS enzyme can be activated by multiple physiological (oxygen and shear stress) (Hickok et al., 2013) or humoral (bradykinin and insulin) cues that drive diverse phosphorylation events on multiple sites such as serine 114, 615, 633, 1177, 1179 and threonine 495 in the oxygenase and reductase domains of the protein (Bauer et al., 2003). When released from the endothelium, NO diffuses into vascular stromal cells such as pericytes or smooth muscle cells to promote vascular relaxation through the stimulation of cyclic guanosine monophosphate (cGMP synthesis by soluble guanylyl cyclase (Archer et al., 1994; Murad, 2004; Russo et al., 2002).

Dysregulation of NO signaling pathways is associated with pathophysiological conditions such as endothelial dysfunction, a main driver for atherogenesis and cardio-metabolic disorders (Kingwell, 2000; Pechánová et al., 2015; Trochu et al., 2000). In endothelial dysfunction, reduced availability of NO converts the endothelial cell phenotype to a pro-inflammatory state with increased oxidative stress and a loss of vasodilatory capacity (Feng and Hedner, 1990; Flammer and Lüscher, 2010). At the molecular level, multiple factors have been reported to contribute to the loss of NO generation in endothelial dysfunction, including a deficiency in the substrate L-arginine or co-factors such as flow-mediated dilation (FMD), nicotinamide adenine dinucleotide phosphate (NADPH), or tetrahydrobiopterin (BH_4_). For instance, when BH_4_ is converted to BH_2_ (7,8-dihydrobiopterin) in an oxidative environment, the eNOS enzyme gets uncoupled from the cofactor leading to the generation of superoxide rather than NO (Bevers et al., 2006; Chen et al., 2010; De Pascali et al., 2014; McNeill and Channon, 2012; Wu et al., 2014; Yang et al., 2009).

At the physiological level, it is well established that fluid shear stress enhances the activity of endothelial nitric oxide synthase (Cabral et al., 2010; Tanaka et al., 2021). Endothelial cells detect changes in local hemodynamics through mechanosensors located on the luminal side of the membrane, in the focal adhesions in the abluminal side, and in the cell-cell junctional complexes. When exposed to laminar shear stress, endothelial cells activate and mobilize Ca^2+^ from intercellular stores. This mobilization leads to the formation of Ca^2+^/calmodulin complexes, which then bind and activate the eNOS enzyme (Corson et al., 1996). Furthermore, fluid shear stress induces the eNOS activation through the tyrosine phosphorylation of PECAM-1 (platelet endothelial cell adhesion molecule-1). This process enhances the phosphorylation of eNOS at Ser^1177^ and Akt Ser^473^ residues. In contrast, lower levels of activation are observed in static conditions in human umbilical cord vein endothelial cells (HUVECs) (Fleming et al., 2005).

Under laminar flow conditions, endothelial cells exhibit an atheroprotective phenotype. However, when laminar flow is disturbed or absent, the cells shift to an inflamed phenotype characterized by the activation of NF-κB (Nagel et al., 1999). This shift in phenotype can contribute to thermogenesis, inflammation, atherosclerosis (Tovar-Lopez et al., 2019; Williams et al., 2021) and obstructive pulmonary disease (Barak et al., 2017). Given the crucial role of shear stress in accurately modeling physiological endothelial responses to risk factors, it is essential for *in vitro* models to incorporate environmental cues such as laminar shear. This necessity is underscored by studies indicating that static endothelial cell-culture models exhibit a proinflammatory phenotype (Junaid et al., 2020), display features of complement-mediated injury (Cabrera et al., 2022) and show impaired nitric oxide (NO) production (“Between Rho(k) and a Hard Place | Circulation Research,” n.d.; Schaefer and Hordijk, 2015). When it comes to developing 3D models, microfluidics emerges as an optimal technology for introducing flow into a 3D microvascular structure (Junaid et al., 2020; van Duinen et al., 2017). The accurate measurement of NO poses a challenge due to its strong reactivity, short half-life, and low physiological concentrations. Traditional techniques such as the Griess assay, chemiluminescence, and amperometric NO sensor readings face limitations in accuracy and sensitivity (Hunter and Schoenfisch, 2015; Moon et al., 2016; Sandrini et al., 2010; Taylor et al., 2006; Vidanapathirana et al., 2019). However, measuring stable metabolites involved in NO production provides a viable approach to quantify NO, even at the low abundance in microfluidic cell models.

In particular, tracer-based metabolomics stands out as the gold standard method. This approach utilizes stable isotope-labelled precursors to trace complex pathways by following the labeled atom(s) to downstream metabolites (Chokkathukalam et al., 2014; Paul Lee et al., 2010). This technique, applied to enzymatic reactions involved in NO production (Figueroa et al., 2020; Gambardella et al., 2020; Kucharzewska et al., 2010; Shatanawi et al., 2020; Shin et al., 2015; Siervo et al., 2011) enhances accuracy and sensitivity, overcoming the challenges associated with traditional NO measurement methods.

To evaluate the influence of adverse metabolic and inflammatory plasma factors on eNOS activity in both 2D and 3D endothelial cell models, we aimed to employ our tracer-based metabolomics strategy. In our study, we used ^13^C_6_,^15^N_4_ L-arginine as a substrate for eNOS, and traced its downstream metabolites ^13^C_6_,^15^N_3_ L-citrulline (a co-product of NO production produced at equal molar amounts) and ^13^C_5_,^15^N_2_ L-ornithine (contributing to citrulline production).

To assess these L-arginine downstream metabolites, we employed an optimized AccQ-Tag amino acid derivatization sample preparation method coupled with a targeted LCMS (liquid chromatography and Mass spectrometry) approach. To validate the accuracy of our method in reflecting endothelial cell biochemistry, we measured changes in metabolite flux in cell models treated with stimulatory and inhibitory compounds for eNOS and arginase. Furthermore, we investigated the impact of unidirectional fluid flow on metabolic parameters associated with NO synthesis. This involved a comparison between a three-dimensional (3D) model with controlled fluid flow and a 2D static cell culture model. The results obtained highlight the suitability of the LCMS method for measuring low-volume and low-abundance NO downstream metabolites. Notably, the 3D model reveals significant effects in both control and VEGF stimulation, as evidenced by changes in Citrulline+9/Ornithine+7. In contrast, static culture lacks significance, indicating a non-regulated eNOS and arginase pathway, resulting in a ED condition. Furthermore, under flow conditions, there are elevated levels of mechanosensitive gene expression compared to static, low shear and bidirectional flow statuses. These results underscore the potential importance of shear stress and NO metabolite levels in influencing endothelial responses and gene expression patterns.

## 2. Material and Methods

### 2.1. Chemicals and reagents

Vascular Endothelial Growth Factor (VEGF) (PeproTech 450-32), L-NAME (abcam, ab120136), BEC (Sigma Aldrich, 63107-40-4), BH4 (Merck, 69056-38-8). RPMI SILAC - medium deficient in Arginine was obtained from (Thermofisher, 88365). Isotope labelled L-Arginine was obtained from (CORTECNET, 130541). Cells were grown under 5% CO2, with 95% atmospheric air. Oxygen experiments were performed in a Panasonic 97 oxygen incubator. DAF-2D was obtained from (Abcam, ab145283).

### 2.2. Cell culture and experimental procedures

Human coronary artery endothelial cells (HCAECs) (PomoCell, C-12221,) were resuspended in 10 ml fresh EGM MV2 medium with supplements (PromoCell,C-22022, C-39216) and cultured in T75 flasks (Nunc Easyflask, Sigma, F7552). Cell cultures were maintained at 37 ^0^C with 5% CO_2_ and media was refreshed three times a week. Cells were detached at 85% confluence with 0.25 % Trypsin EDTA (Lonza, CC-5012) and cell pellets were collected by centrifugation at 300g for 5 minutes.

For 2D culture experiments, the collected cell pellets were suspended in a fresh medium to a concentration of 7 x 10^5^ cells/ml and cultured in a 48 well plate for 48 hours. The following day, cells were serum starved with 1% FCS in basal EGM2 (Bioconnect, C-22216) medium and incubated overnight. To synchronize or equilibrate the cell cycle we treated cells with Krebs buffer solution, pH 7.4 (Thermo fisher Scientific, J67795.AP) for 1 hour before the start of each experiment. Subsequently the cells were incubated for 12 hours with RPMI SILAC (ThermoFisher Scientific, 88365) supplemented with 150µM ^13^C_6_, ^15^N_4_-L-arginine (CORTECNET, 130541), 10 µM of (6R)-5,6,7,8,- Tetrahydrobiopterin dihydrochloride (BH_4_) (Merck, 69056-38-8) and 1% FBS (Thermofisher Scientific, A4736301) in the presence of 100ng/ml VEGF (PeproTech 450-32), 1mM L-NAME (MedChemExpress, HY-18729A) or 100µM BEC (Sigmaaldrich, SML1384) separate or in the indicated combinations. The medium and cells were collected separately and snap freeze in liquid nitrogen and stored in -80°C for LCMS analysis.

For 3D cultures, we used a modified chip design 2-lane rerouted OrganoPlate (MIMETAS, Netherlands) that was adapted to attach a microfluidic pump to apply fluid flow and normal 2-lane OrganoPlate (9603-400-B) to apply bidirectional flow. The incubation protocols were similar to that of the 2D assay described above. After seeding HCAECs cells into the 3D microvessels, they were cultured for 3-4 days with media refreshed every other day. The microvessels were treated with VEGF, L-NAME and BEC as described above for 12 hours. Following the application of shear stress and compound treatments, media samples were collected, snap freeze with liquid nitrogen and stored at -80°C for LCMS analysis.

### 2.3. Immunofluorescence microscopy

HCAECs were fixed using 4% paraformaldehyde (PFA) in HBSS+ for 10 min at room temperature. The fixative was aspirated, and the cells were rinsed once with HBSS+. The cells were permeabilized for 2 min with 0.2% Triton X-100 in HBSS+ and washed with HBSS+. Permeabilization was followed by blocking cells using 5% BSA in HBSS+ for 30 min and incubated with the primary antibody solution - Mouse anti-human CD144 (1:150; 555661, BD Biosciences, USA) for overnight at 4°C. The cells were washed with HBSS+, followed by a one-hour incubation with Hoechst (1:2000; H3569, Invitrogen, USA), rhodamine phalloidin (1:200; P1951, Sigma-Aldrich, The Netherlands) and the secondary antibody solution, containing Alexa Fluor 488-conjugated goat anti-mouse (1:250; R37120, ThermoFisher, USA). The cells were washed three times with HBSS+. High-quality images of the stained cells were acquired using a high-content confocal microscope (Molecular Devices, ImageXpress Micro Confocal).

### 2.4. Sample preparation for LC-MS

Samples were analyzed using a method based on amine profiling platform that employed an AccQ-Tag derivatization strategy, adapted from the protocol provided by Waters (Noga et al., 2012). The cell-treated media and cell lysates (10 µL) were thawed on ice. For deproteination, water (5µL), Tris-(2- carboxyethyl) phosphine hydrochloride (TCEP) (10 µL), and absolute methanol (75 µL) were added. Quality control (QC) samples were generated by pooling equal volumes of all cell media and cell lysate samples, and 10 µL of this pool underwent the same process as individual samples. After centrifugation at 13,200 rpm for 10 minutes, the supernatant was transferred to a fresh Eppendorf tube and dried under speed vacuum. The dried residue was reconstituted in borate buffer – pH 9 (10 µL) vortexed for 10s and treated with 2.5 µL of AccQ-TagAQC derivatization reagent (Waters, Waters B.V. Art. No. 186003836, The Netherlands). The samples were then kept at 55^°^C for 30 minutes in a shaker (Incubating microplate shaker, VWR, The Netherlands), 20% of formic acid (5 µL) was added for neutralization. After a quick vortex, each sample was transferred to a deactivated auto sampler vial for LC-MS injection.

### 2.5. Instrumentation and LC-MS acquisition

Three µl of sample solution was injected onto a UPLC Class I (Acquity, Waters Chromatography Europe BV, Etten-Leur, The Netherlands) system with an AccQ-Tag Ultra C18 Column (1.7 µm, 100 x 2.1 mm, Waters, Ireland) coupled to a Sciex QTRAP® 6500 mass spectrometer (Noga et al., 2012). For liquid chromatography (LC) separation, mobile phase A consisted of 0.1% formic acid in water. Mobile phase B consisted of 0.1% formic acid in acetonitrile. The flow rate used was 0.7 mL/min and the starting gradient condition was 99.8% A for 0.5 min, changing linearly to 90% A over the next 5.50 min, 80% A over 7.50 min, 40% A at 8.00 min, 5.0% A at 9.00 min, after which the solvent composition returned to 99.8% A over next 9.10 min and ends up with 99.8 % A and 0.2% B over in next 11.00 min. Mass spectrometry experiments were carried out on a SCIEX QTRAP 6500. The ESI source parameters were as follows (positive ion mode): Spray voltage ± 5.5kV, capillary temperature 350°C, sheath gas 60 psi, auxillary gas 70 psi, curtain gas 30 psi. Data acquisition was performed in multiple reaction monitoring mode targeting compounds with different labeling statuses. The compound list with target *m/z* for parent and product ions is shown in **Supplementary Table. 1.** Raw LC-MS/MS data was processed with AB Sciex PeakView™ 2.0 and MultiQuant™ 3.0.1 software for targeted metabolite peak identification and integration. Metabolite isotopologue levels were quantified using the corresponding peak areas.

### 2.6. Microfluidic pump setup

HCAECs were seeded into an OrganoPlate (similar protocol mentioned in the above section) that was specially designed for controlled perfusion using a pump attachment. The pump is made of stainless steel and the tube connections for the fluid transfer from inlet and outlet of the chips are made of silicone. For controlled rotation, a small motor is operated with the help of LabVIEW solutions software. This software allows the control of fluid flow rate and related shear stress (dyne/cm^2^), use of portal connections, motor position and value. Typically, shear stress was applied for 12 hours for metabolic readouts and 12 and 24 hours for gene expression studies to condition the cells.

### 2.7. Validation of eNOS activity using DAF-2D

HCAECs were cultured in a 48 well plate with 1.5 × 10^3^ cells/ well suspended in a fresh EGM MV2 medium with supplements. Upon 80% confluence, the cells were starved with 1% FCS in EGM basal medium for 17 hours at 37°C with 5% CO_2_. Next the cells were treated with Krebs buffer solution for 1 hour at 37^°^C. After two washes of PBS, 5µM of DAF-2D stain (Abcam, ab145283) was added and the cells were incubated for 1 hour at 37°C and then washed twice with PBS. The cells were treated with VEGF, L-NAME, BEC and only the working solution (basal EGM2 media with 1% FCS and 10µM BH_4_) separately and incubated for 10-12 hours as described before. The cells were washed twice with PBS and imaged in EVOS CO_2_ incubator build fluorescent microscope with maximum excitation: 491nm and maximum emission: 513nm. The fluorescence intensities were measured by choosing ten random individual cells and grey scale intensity was analyzed using imageJ software to correlate with citrulline level.

### 2.8. Quantitative RT-PCR

RNA was extracted using Qiagen’s RNAeasy kit and Buffer RLT lysis buffer based on the manufacturer’s recommendations. Then, to enable analysis of gene expressions at low concentrations due to low cell counts, the lysate of eight microvessels from 3D cell culture models and eight well samples from 2D -48 well plate were merged to make one sample. Reverse transcription-mediated cDNA synthesis was carried out using random and oligo(dT) primers (Bio-Rad) in accordance with the manufacturer’s instructions using 200 ng of total RNA. For the qRT-PCR analysis, SYBR Select (Invitrogen) and a Biorad CFX384 were utilized. Klf2 (sense), CTACACCAAGAGTTCGCATCTG; Klf2 (antisense), AGCACGAACTTGCCCATCA; 18S rRNA (sense), GGATGTAAAGGATGGAAAATACA; 18S rRNA (antisense), TCCAGGTCTTCACGGAGCTTGTT were the target genes whose primer sequences were employed. Expression levels were standardized to 18S rRNA and quantified using the comparative cycle threshold (ΔΔCt) method.

### 2.9. Statistical analysis

Bar plots and box plots were created with GraphPad Prism 9.3.1 software. Significance determined by ANOVA, Tukey’s analysis and student t-test.

## 3. Results and Discussion

### 3.1. Tracer-based measurement of eNOS-dependent arginine to citrulline conversion - Marker metabolites that reflect eNOS activity

The purpose of this study was to understand endothelial NO production from a metabolic standpoint. We examined isotope-labeled arginine-based NO metabolic modifications by employing specific stimulant and inhibitory compounds targeting eNOS and arginase enzymes. Our focus was on measuring the impact of adverse metabolic and inflammatory factors on NO synthesis in endothelial cells, and for this purpose, we developed a highly sensitive tracer-based method to quantify eNOS-dependent conversion of arginine to citrulline and ornithine.

The LCMS method utilized in this study is targeted for the analysis of isotope-labeled L-arginine and its downstream metabolites, L-Citrulline and L-Ornithine, using AccQ-Tag derivatization. The metabolic pathway of L-arginine metabolism and its conversion to downstream metabolites, as illustrated in (**Fig. 1A),** was the focal point of our analysis. We optimized the method by analyzing potential isotopologues of these marker metabolites in the cell system **(Suppl. Table. 1)**.

**Figure. 1.**
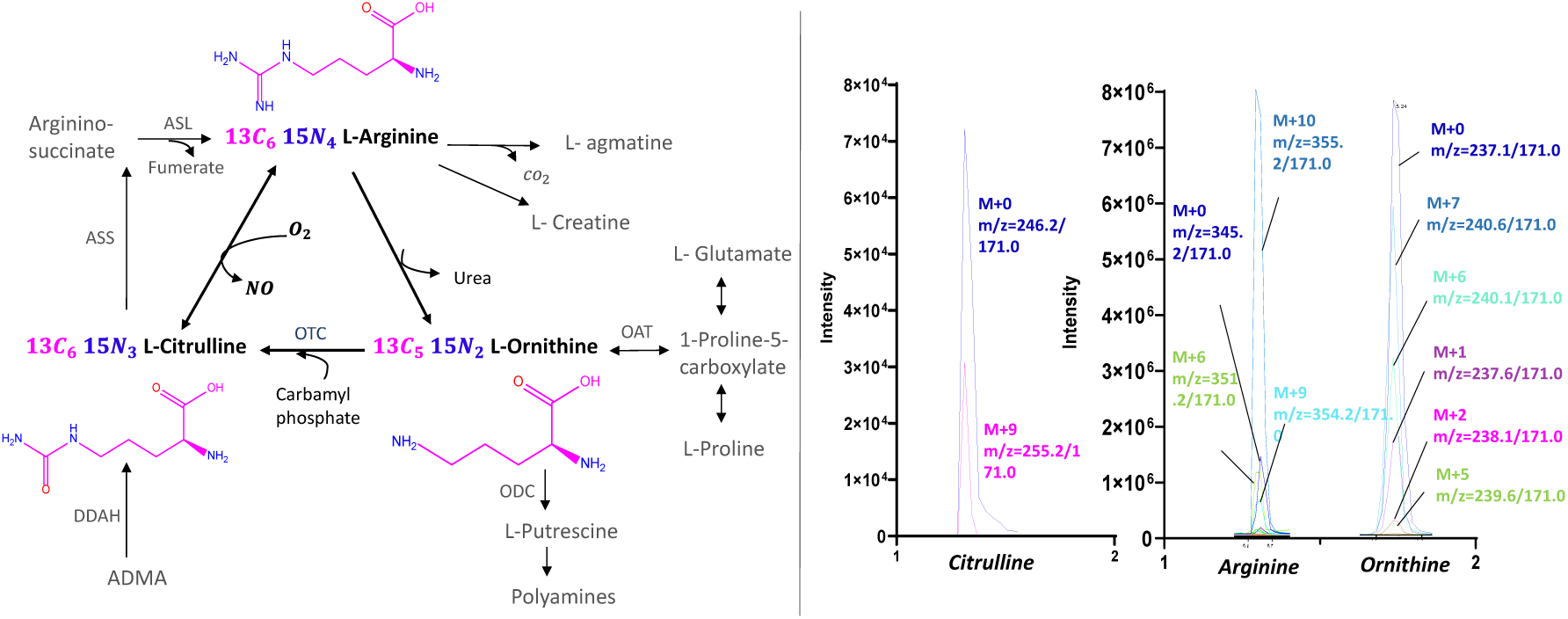
Tracer-based Nitric Oxide (NO) metabolomics study. (A) Metabolic pathway of L-arginine and a few metabolites involved in other functions. Three marker metabolites (in bold) were analyzed based on stimulated and inhibited conditions of the respective enzymes eNOS and arginase. (B) Representative ion chromatograms of metabolite isotopologues detected in HCAECs sample obtained after cells were incubated with 150 µM of, ^13^C_6_ ^15^N_4_ L-arginine.

After treating the human coronary artery endothelial cells (HCAECs) with isotope labelled L-arginine, we analyzed the resulting samples to identify peaks representing a series of Isotopomeric citrulline, arginine and ornithine in the ion chromatogram **(Fig. 1B).** Arginine was detected with M+10 (representing ^13^C_6_, ^15^N_4_ labeling) as a primary isotopologue, accounting for over 90% in all samples. Citrulline was detected with M+9 (representing ^13^C_6_, ^15^N_3_) as the primary isotopologue, produced by losing one nitrogen. Ornithine was detected with M+7 (representing ^13^C_5_, ^15^N_2_), resulting from the loss of two carbons and two nitrogen atoms from arginine. In the subsequent analysis, ^13^C,^15^N-arginine (M+10), ^13^C,^15^N-citrulline (M+9) and ^13^C,^15^N-ornithine (M+7) were selected as biomarkers for further investigation. The measurements of NO metabolites, free from non-metabolism-related artefact signals as background, were compared with the blank values (only media). This comparison confirmed there was no high background signal of the downstream metabolites (results not shown).

To optimize the incubation time, we considered the effects observed with stimulatory and inhibitory compounds. We utilized the detected isotopologues to verify the uptake of ^13^C_6_, ^15^N_4_-L-arginine and reconstruct the metabolic pathways controlled by eNOS and arginase in HCAECs. This analysis encompassed four conditions: 1) control, 2) treatment with the stimulatory compound VEGF, known for inducing NO release from endothelial cells (Feliers et al., 2005); 3) treatment with the eNOS inhibitory compound L-NAME (Rees et al., 1990; Silva et al., 2022); and 4) treatment with the arginase enzyme inhibitor BEC (Caldwell et al., 2015). The pathway and resulting metabolite expression for each treatment are depicted in (**Fig. 2 A-C)**. The optimization process led us to select a 12-hour incubation time after the introduction of tracers. We observed lower labeling incorporation at 3 and 6 hours, as indicated by the ratio of Citrulline+9/Arginine+10 and Ornithine+7/Arginine+10 **(Suppl. Fig. 1).** This rate of label incorporation could be influenced by the exchange of internal and exterior metabolites. Thus, we opted for a 12-hour treatment to see the maximal transfer of atoms. The calculated isotopologue fractions after 12 hours of incubation time shows that the utilization of labelled arginine converted into maximum transition state of labelled Citrulline (M+9) and ornithine (M+7) **(Suppl. Fig. 2)**. Despite the brief treatment time, the obtained marker metabolites were relatively stable and high after 12 hours, therefore, further analyses were performed on the samples produced after 12 hours of incubation.

**Figure. 2.**
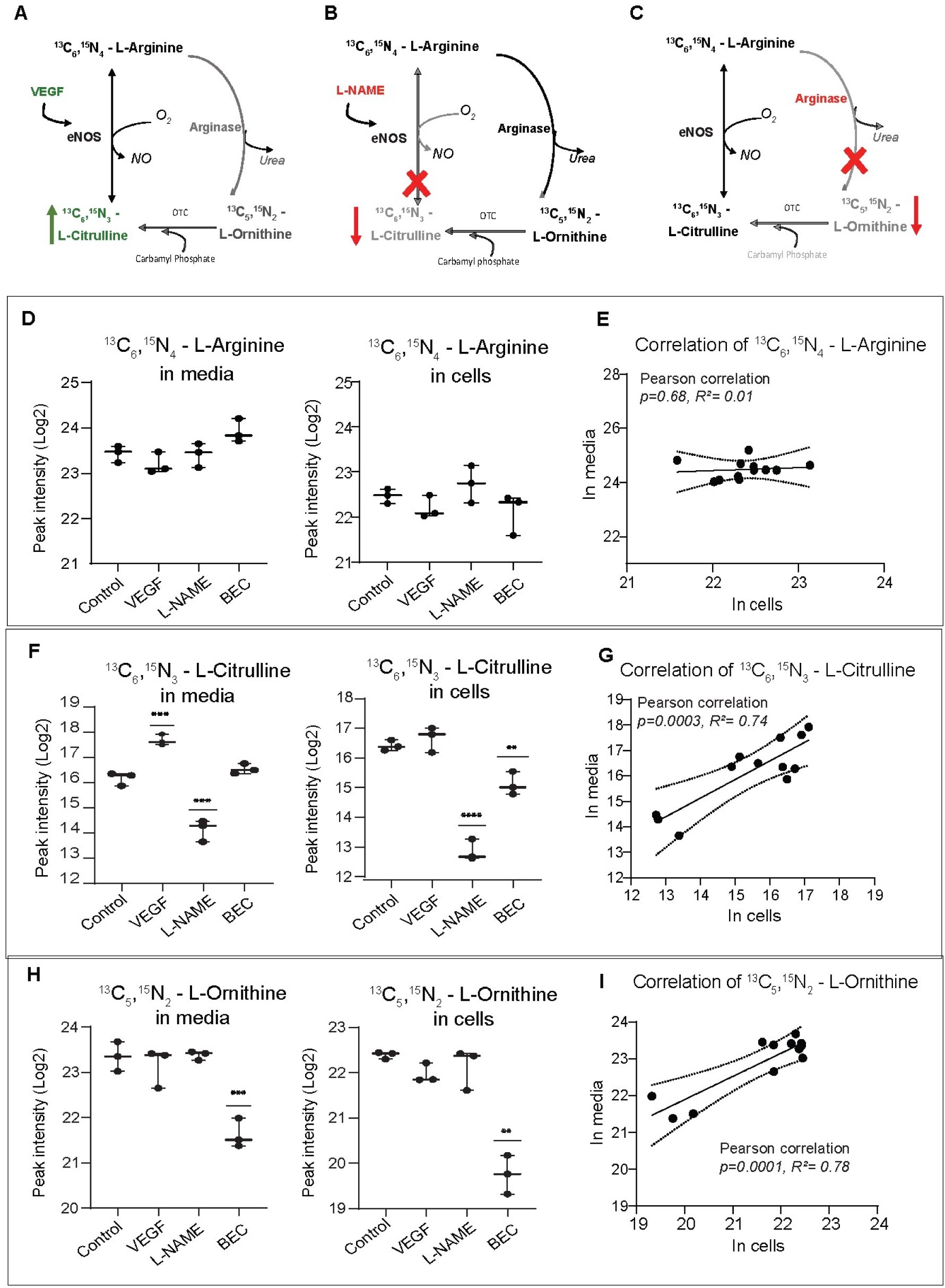
Measurement of isotope labelled extracellular (media) and intracellular (cells) marker metabolites involved in NO mechanism. A) Schematic representation of NO pathway was highlighted upon treatment of A) VEGF – eNOS stimulation (green) B) L-NAME - eNOS inhibition (red) and C) BEC – arginase inhibition (red). Highlighted and non-highlighted paths are the representation of active and inhibition conditions. Measurement of metabolites in media and cells (D) ^13^C_6_,^15^N_4_ L-arginine and its (E) Pearson correlation; (F) ^13^C_6_,^15^N_3_ L-Citrulline and its (G) Pearson correlation; (H) ^13^C_5_,^15^N_2_ L-Ornithine and its (I) Pearson correlation were mentioned respectively. All error bars represents SD and mean, n=3. Significance determined by student t-test of treated group versus control group. *,P<0.05; **,P<0.01; ***,P <0.001; ****,P<0,0001.

### 3.2. Extracellular and intracellular measurement of Isotopomeric compounds

In order to comprehensively understand the effects of stimulatory and inhibitory compound treatments and to evaluate cellular metabolism, we conducted measurements of marker metabolites both intracellularly and extracellularly in a 2D cell culture platform. HCAECs were treated with eNOS stimulator (VEGF), eNOS inhibitor (L-NAME) and arginase inhibitor (BEC) in conditioned media containing ^13^C_6_,^15^N_4_ L-arginine for 12 hours. Subsequently, both extracellular (media) and intracellular (cell) samples were analyzed in LC-MS/MS. The targeted metabolic routes for individual treatments are illustrated in (**Fig. 2 A-C)**.

In examining the results of specific treatments, we observed that ^13^C_6_,^15^N_4_ L-arginine exhibited lower intracellular levels compared to extracellular secretions in all conditions **(Fig. 2D)**. Statistical analysis indicated a non-significant correlation between media and cells for ^13^C_6_,^15^N_4_ L-Arginine (Pearson correlation coefficient *P value*= 0.6, R^2^ = 0.01) **(Fig. 2E).**

The level of ^13^C_6_,^15^N_3_ L-Citrulline was higher in media than within cells **(Fig. 2F).** In earlier studies, VEGF stimulation was shown to enhance the NO production through the activation of tyrosine kinase activity, resulting in the phosphorylation of intracellular domains. This activation increases calcium levels, facilitates binding with calmodulin (CaM), enhances eNOS phosphorylation, and ultimately leads to increase NO production (Grover et al., 2002; Pandey et al., 2018; Cazzaniga et al., 2018; Gélinas et al., 2002). The mechanism via receptor activation is depicted in **Suppl. Fig. 3**. Our results indicate that eNOS activation leads to increased citrulline, consistent with its role as a co-product of NO production. The eNOS inhibition with L-NAME treatment shows significantly lower ^13^C_6_,^15^N_3_ L-Citrulline and in arginase inhibition the level is lower in cells than in media. Overall, our result show that there is a significant positive correlation between media and cells for ^13^C_6_,^15^N_3_ L-Citrulline (Pearson correlation coefficient, p = 0.0003, R^2^ = 0.7) indicating a significant positive connection **(Fig. 2G)**. The ^13^C_5_,^15^N_2_ L-Ornithine levels were not significantly affected by VEGF and L-NAME treatments, in contrast to the condition where a specific inhibition of the arginase enzyme with BEC treatment was added, resulting in decreased ornithine levels **(Fig. 2H).** When we measured ^13^C_5_,^15^N_2_ L-Ornithine in media and cells, a significant positive correlation with *p* value = 0.0001, R^2^ = 0.7 was observed **(Fig. 2I)**. Earlier studies have reported BEC as a classical competitive inhibitor of arginase II at pH 7.5 with a dissociation constant (Ki) describing the binding affinity between the inhibitor and the enzyme as 0.25 and 0.31µM (Colleluori and Ash, 2001). Therefore, both citrulline and ornithine, upon stimulatory and inhibitory compounds treatment, exhibited similar effects between media and cells. In general, most of the *in vitro* analyses measured NO metabolites only in intracellular level (Hecker et al., 1990; Tsuboi et al., 2018). In our study, we attempted and proved that the obtained promising results showed extracellular and intracellular expressions are positively correlated and this study is helpful for fluxomics study. As a result of this study, our further analysis were done in extracellular samples.

### 3.3. Analysis of metabolic ratios to understand NO metabolic flux

Metabolite ratios can serve as valuable indicators of diseased conditions, providing insights into the underlying biological mechanisms and disrupted pathways in the disease state (Molnos et al., 2018). Using the extracellular measurement method mentioned earlier, our focus was on ratios that reflect metabolic fluxes, and we conducted investigations using individual and combined treatments of endothelial cells with stimulatory and inhibitory compounds targeting eNOS and arginase enzymes. This approach allowed us to gain a comprehensive understanding of how these compounds, both individually and in combination, influence the metabolic pathways associated with endothelial dysfunction.

Under eNOS stimulation conditions (VEGF, Pathway **Fig 2A**) the expected outcome is a maximized production of citrulline compared to conditions involving arginase and eNOS inhibition (Pathway **Fig 2B & C**). The ratio ^13^C_6_,^15^N_3_ L-Citrulline/ ^13^C_6_,^15^N_4_ L-arginine was found to be significantly higher in the VEGF treatments, indicating a greater utilization of arginine for citrulline conversion (**Fig. 3A**). This ratio has been extensively examined in children with chronic kidney disease (CKD) and abnormal cardiac blood pressure conditions, reporting a higher citrulline-to-arginine (cit-arg) ratio in plasma (Lin et al., 2013) but a lower ratio reported in urine (Lin et al., 2016). This research indicates that evaluating the cit-arg ratio proved to be an effective predictor of cardiovascular prognosis in children and adolescents with early CKD.

**Figure. 3.**
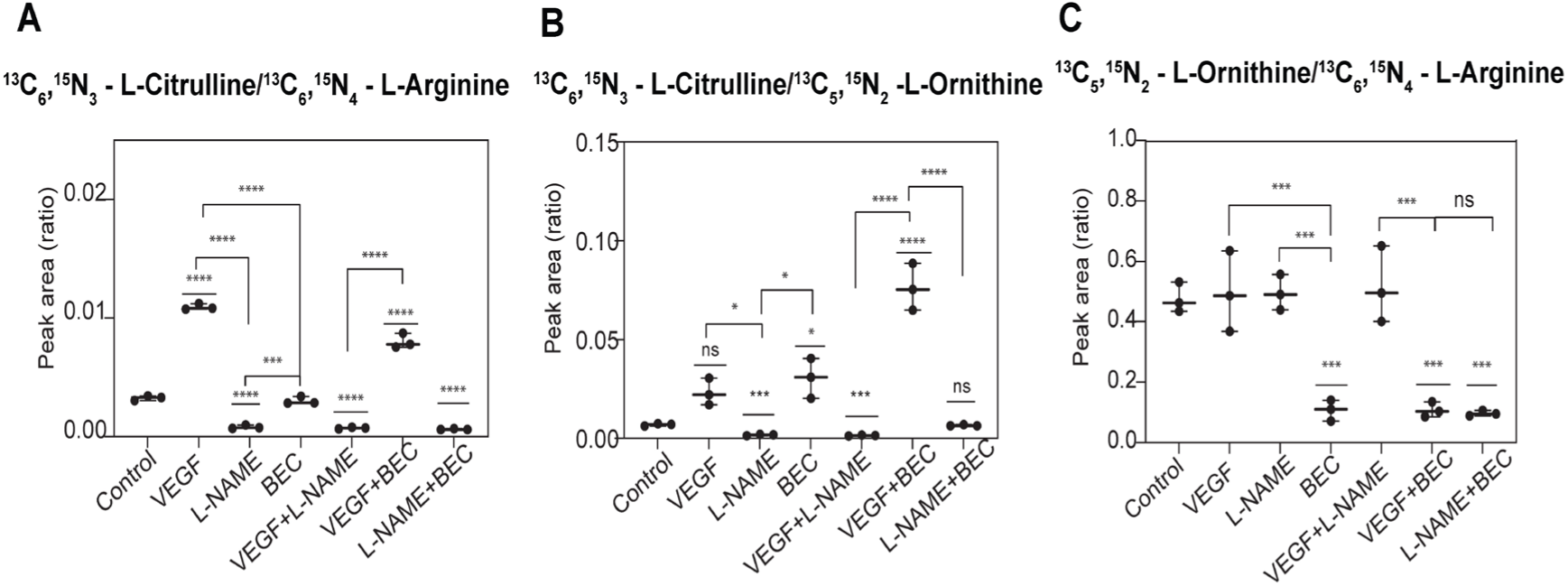
Measurement of extracellular labelled metabolites in 2D platform with the treatment of stimulators and inhibitors in an individual and combinational treatments. (A) Ratio of ^13^C_6_,^15^N_3_ L- Citrulline to ^13^C_6_ ^15^N_4_ L-arginine (B) ratio of ^13^C_6_,^15^N_3_ L-Citrulline to ^13^C_5_,^15^N_2_ L-Ornithine and (C) ratio of ^13^C_5_,^15^N_2_ L-Ornithine to ^13^C_6_ ^15^N_4_ L-arginine. All error bars represent SD and mean, each dot represents biological replicates. Significance determined by ANOVA and Tukey’s multiple comparison test. ns = not significant; *, P<0.05; **, P<0.01; ***, P <0.001; ****, P<0,0001.

The ^13^C_6_,^15^N_3_ L-Citrulline /^13^C_5_, ^15^N_2_ L-Ornithine ratio serves as an indicator of the balance in the metabolic pathway associated with arginase and eNOS activities. Our result shows a significant increase is observed in arginase inhibition conditions (BEC, and VEGF+BEC) **(Fig 3B).** The higher ratio resulting from the BEC treatment suggests that arginase inhibition could be a potential target for recovering NO or citrulline production in diseased conditions. The inhibition of eNOS by L-NAME treatments (L-NAME, and VEGF+L-NAME) **(Fig.3B)**, led to lower levels of citrulline and ^13^C_6_,^15^N_3_ L-Citrulline /^13^C_5_, ^15^N_2_ L-Ornithine ratio indicating that the eNOS inhibition was selective.

The ^13^C_5_,^15^N_2_ L-Ornithine/^13^C_6_,^15^N_4_ L-arginine ratio, analyzed to confirm the utilization of arginine to ornithine conversion, showed lower expression in all arginase inhibition (BEC) conditions compared to other treatments **(Fig 3C).** Our data supports the idea that the arginase inhibition recovers citrulline levels better than the stimulated condition with more arginine availability (Shatanawi and Momani, 2018).

Furthermore, the combinational treatments provide insights to the metabolic alterations in a controlled fashion. For instance, in VEGF+BEC condition, production of ^13^C_6_,^15^N_3_ L-Citrulline was directly from arginine rather than from ornithine as the pathway is blocked by the arginase inhibitor BEC. Such information is valuable for comparing the underlying mechanisms of disease and identifying potential targets for therapeutic interventions.

### 3.4. Validation of eNOS activity using NO specific DAF-2DA staining

To determine if citrulline production can serve as an estimate eNOS activity, we compared the peak intensities of ^13^C_6_ ^15^N_3_ L-citrulline with another measure of eNOS activity - Diaminofluorescein-2 diacetate (DAF-2DA) fluorescence staining for detecting intracellular NO **(Fig. 4)**.

**Figure. 4.**
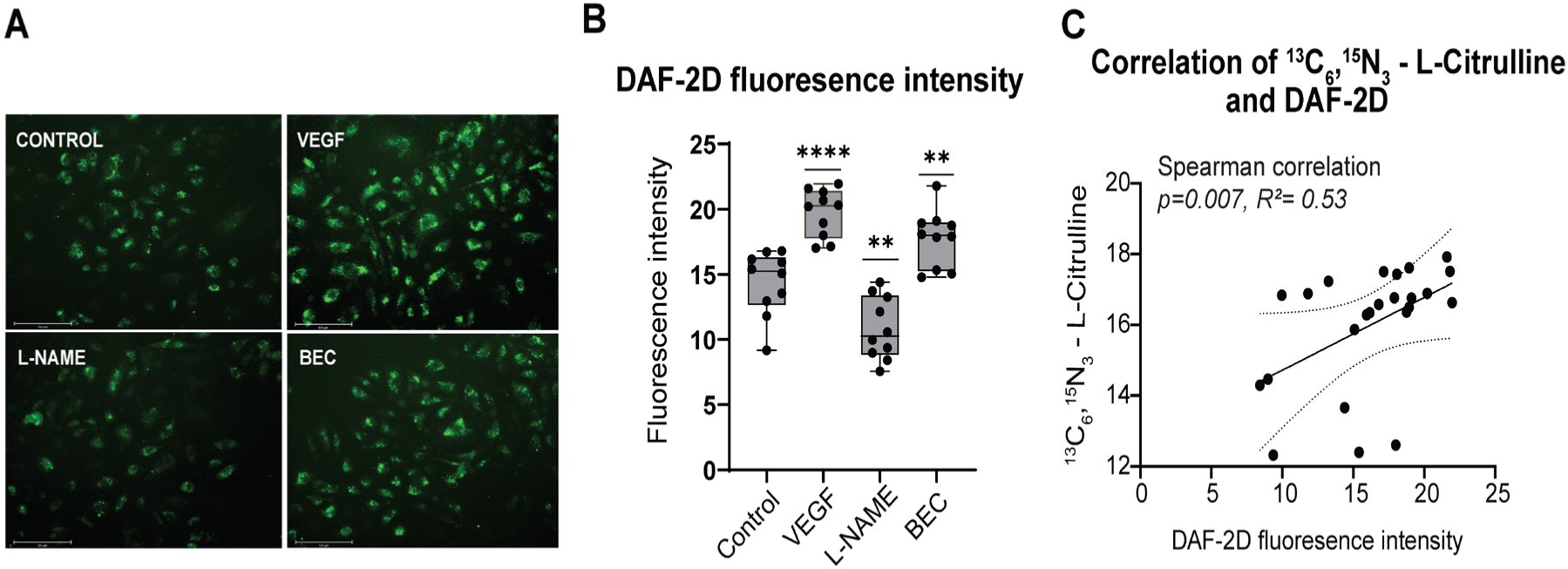
NO specific DAF-2 DA fluorescent stain on HCAECs. (A) Microscopic images of HCAECs using fluorescent NO stain DAF-2DA in all treatments; eNOS stimulator (VEGF), inhibitor (L-NAME) and arginase inhibitor (BEC). (B) Intensity of the fluorescent stain was calculated by grey scale measurement using ImageJ software; n=10. (C) Spearman correlation of ^13^C_6_,^15^N_3_ L-Citrulline and DAF-2DA fluorescence intensity. Significance determined by student t-test. ns = not significant; *, P<0.05; **, P<0.01; ***, P <0.001; ****, P<0,0001.

This experiment was conducted in the 2D platform (48 wells cell culture plate) under the same conditions used for metabolic read out, followed the technique as described in the preceding section. DAF-2D fluorescence intensity is significantly higher in the VEGF condition, indicating an enhancement of NO production, as previously demonstrated in HUVECs cells (Jo et al., 2017). This is consistent with the observations that VEGF stimulation increases both ^13^C_6_,^15^N_3_ L-Citrulline levels and DAF-2D signals in HCAECs **(Fig. 4 A & B)**. Similarly, increased DAF-2D intensity was observed in BEC treatment, as reported earlier in human aortic endothelial cells (HAECs) where arginase inhibition restored NO production (Ryoo et al., 2008), and inhibited NO production in L-NAME treatment **(Fig. 4 A & B).**

To compare the relation between NO level and citrulline production, we performed the correlation analysis. Overall, the statistical analyses of the data indicate significant correlation between DAF-2D fluorescent intensity **(Fig. 4B)** vs. ^13^C_6_ ^15^N_3_ L-citrulline level (Spearman correlation coefficient, p = 0.007, R^2^ = 0.53) denotes that the amount of ^13^C_6_ ^15^N_3_ L-citrulline production from 2D model **(Fig.2F**) is comparable to the NO level using this staining method **(Fig. 4C).** Although, the direct stain was not successful in our 3D microvessels. The 3D model is shown in the **Fig. 5A-E**. The major disadvantage we encountered was, the dye has given strong background signal when reacting to the collagen which acts as the extracellular matrix in a 3D microvessel model where cells embedded in one side of it (results not shown). Furthermore, major disadvantages were reported in earlier studies that i) fluorescein could easily photobleached by intense excitation light, and ii) detection reagent does not directly react with NO, but rather with the oxidized form of NO such as such as nitrogen dioxide (NO_2_), dinitrogen trioxide (N_2_O_3_), peroxynitrite (ONOO^-^), nitrite (NO_2_ ^-^), nitrate (NO ^-^), and nitroxyl (HNO). This could be a concern in measurements, where the signal should not interfere substantially with NO signal transduction (Kojima et al., 1998). With the aforementioned limitations, our staining procedure in 2D model was inconsistent when we attempted to reproduce the same staining in 3D microvessel model.

**Figure. 5.**
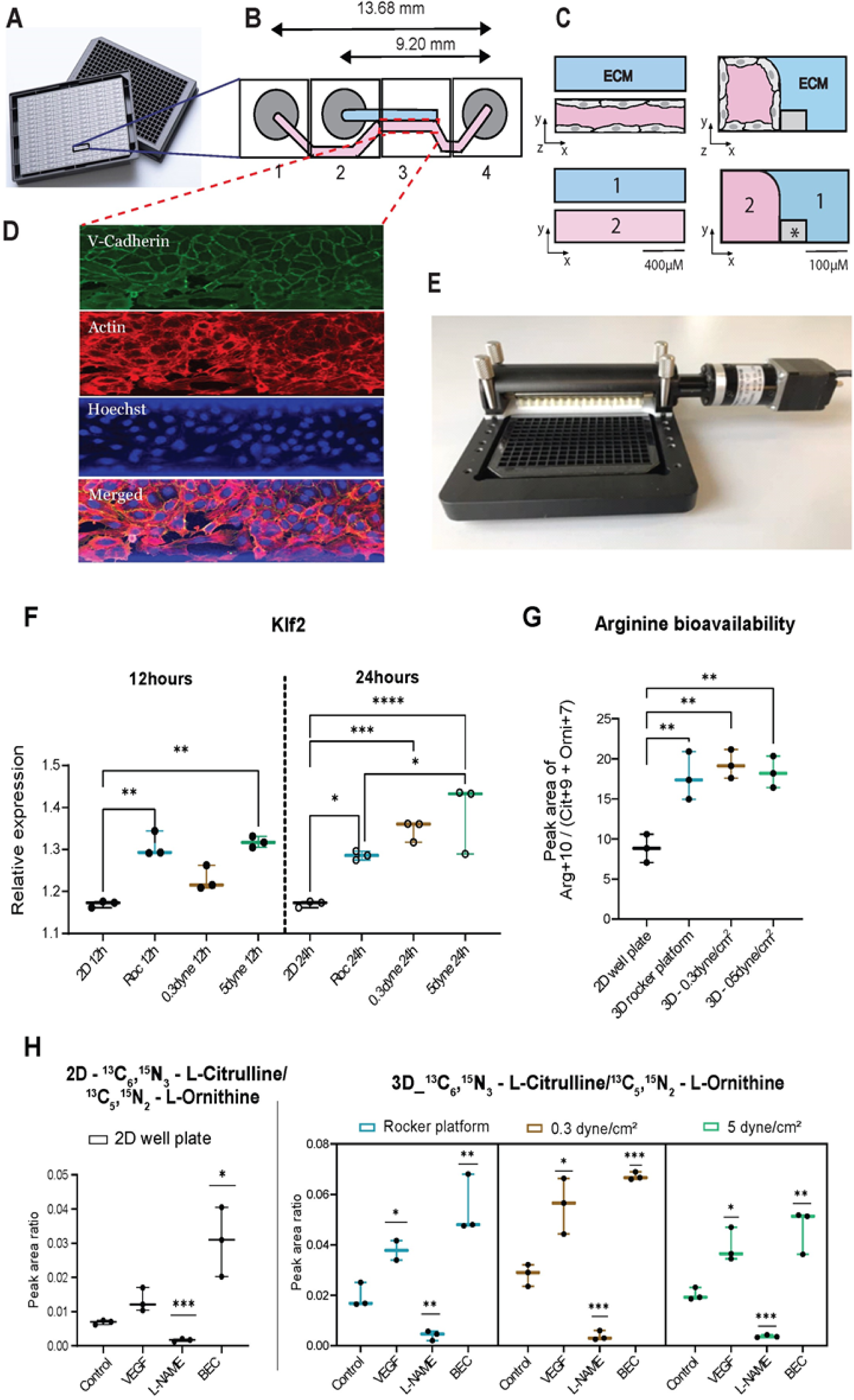
Representation of our 3D blood vessel model and comparison of NO marker metabolites in 2D and 3D platforms. (A) Illustration of re-routed OrganoPlate and its (B) Single chip design was explained below. Every microfluidic chip structure is positioned underneath 4 adjacent wells and it consists of two channels: 1) an ‘perfusion’ channel (pink color) and 2) a ‘gel’ channel (ECM) (blue color). Every first well (1) and fourth well (4) is positioned on top of the inlet and outlet of the perfusion channel, while every second well (2) are for the gel channel. And every third well (3) is used for imaging and observation of the experiment. (C) Perfusion and gel channel was shown vertical and horizontal view and its separated by a phase guide (*). (D) Immunostaining of HCAECs in a chip. After 48 hours of cell seeding, a confluent vessel of HCAECs were immunostained with V-Cadherin, Actin and DAPI. (E) Image of a microfluidic pump with the two metal tubes attached to the inlet and outlet of rerouted OrganoPlate to carry fluid and create unidirectional flow. Ratio analysis and Measurement of extracellular marker metabolites in all platforms. (F) measurement of Klf2 gene expression of cell samples (G) Arginine bioavailability calculation and (H) Ratio of ^13^C_6_,^15^N_3_ L-Citrulline to ^13^C_5_,^15^N_2_ L-Ornithine all obtained from 2D static-black bars), 3D rocker platform (bidirectional flow model – blue bars) & the rerouted OrganoPlate with microfluidic perfusion pump operated with different shear stresses (unidirectional flow model) 0.3 dyne/cm^2^ - brown bars, 5 dyne/cm^2^ - green bars. Data for RTPCR are presented as mean and s.e.m; n = 3. Metabolic data significance determined by one way ANOVA multiple comparison test. *, P<0.05; **, P<0.01; ***, P <0.001; ****, P<0,0001.

### 3.5. Comparison of mechanosensitive Klf2 gene in static (2D) and flow mediated (3D) cell culture models

Flow-mediated vasodilation in endothelium releases the endogenous nitro vasodilator, nitric oxide (Cabral et al., 2010; Cooke et al., 1991; Zhou et al., 2014). To investigate the importance of shear stress on eNOS activation at the gene and metabolic level, we used a 3D culture model utilizing a novel setup – the MIMETAS 2-lane rerouted OrganoPlate **(Fig 5A).**

The commercially available MIMETAS 2-lane OrganoPlate’s chip design was altered (rerouted) by connecting the route of windows 1 and 4 **(Fig. 5B)**, extending the surface area for more cell growth. This design is suitable for fixing a microfluidic pump to create a flow, in contrast to the commercial MIMETAS OrganoPlate, where such a connected route is not present, as it is meant to apply flow using rocker platform that generates bidirectional flow (van Duinen et al., 2017). The microchannels in the OrganoPlate for gel and perfusion (1 & 2) are illustrated in **(Fig. 5C)**. The microchannels in the OrganoPlate were coated with gelatin, preventing HCAECs from growing on glass and enabling them to form stable microvessels which was checked by immunostaining **(Fig 5D)**. The rerouted OrganoPlate was connected to the microfluidic pump is shown in **(Fig. 5E).**

To validate the functionality of our microfluidic pump, we first used quantitative rtPCR to the measure the impact of shear stress on the shear-dependent transcription of the Klf2 gene (Krüppel-like transcription Factor2) (Wang et al., 2006) which also plays an important role in endothelial function, anti-inflammatory, anti-thrombotic and angiogenesis (Zhou et al., 2014; Turpaev, 2020). For this study, we utilized three different cell culture models. They are a) the 2D model in which cells were grown in a static conventional culture, b) a bidirectional flow model – MIMETAS 2-lane OrganoPlate with normal chip design subjected to flow using rocker (van Duinen et al., 2017) and c) a unidirectional flow model - the rerouted plate with microfluidic pump set to two different dyne stresses, namely 0.3, and 5.0 dyne/cm^2^. From these platforms, HCAECs were collected, and RNA was isolated at 12 and 24 hours. The isolated RNA was transcribed to cDNA with the respective primers and quantified using RTPCR technique.

The results of the 12-hour incubation reveal a significant upregulation of the Klf2 gene in microvessels perfused with the microfluidic pump (5 dyne/cm^2^) and the interval rocker and at low shear stress condition (0.3 dyne/cm^2^) the significance level with p value was 0.06 was observed when compared to the 2D static model. Furthermore, the results of the 24-hour incubation demonstrate that cells subjected to 5 dyne/cm^2^ shear stress exhibit a significant upregulation of the Klf2 gene compared to cells in the 2D well plate and microvessels perfused with interval rocker platform **(Fig 5F).** This indicates that our flow model induces substantial upregulation of the shear dependent gene – Klf2, highlighting the effectiveness of our model in providing proper flow conditions to cells and allowing variations in shear stress.

### 3.6. Comparison of NO marker metabolites in static (2D) and flow mediated (3D) cell culture models

Metabolic analysis was performed in the samples obtained from the above mentioned models (in Section 2.5), and ratios of ^13^C_6_,^15^N_3_ L-Citrulline, ^13^C_5_, ^15^N_2_ L-Ornithine and ^13^C_6_,^15^N_4_ L-arginine were evaluated. The advantage of evaluating ratios rather than comparing individual metabolites is that ratios eliminate the need for additional normalization methods such as cell count or protein levels. The ratio of ^13^C_6_ ^15^N_3_ L-citrulline/^13^C_6_,^15^N_4_ L-arginine indicates that the 2D cell culture platform exhibits better citrulline conversion and arginine uptake compared to the unidirectional flow system 3D model. A similar response was seen for the ^13^C_5_, ^15^N_2_ L-Ornithine /^13^C_6_,^15^N_4_ L-arginine ratio **Suppl Fig. 4A**.

The metabolic effects of VEGF, L-NAME and BEC treatments were comparable between 2D and 3D cultures for the above mentioned ratios **Suppl Fig. 4 B & C.** However, the ^13^C_6_ ^15^N_3_ L-citrulline /^13^C_5_, ^15^N_2_ L-Ornithine ratio showed a significant increase in the 3D unidirectional flow condition (0.3 dyne/cm^2^) in control **Suppl Fig. 4 A** and after eNOS stimulation **(Fig. 5H).**

The influence of flow is evident in the ^13^C_6_ ^15^N_3_ L-citrulline /^13^C_5_, ^15^N_2_ L-Ornithine ratio, indicating that flow “regulates” metabolic expression when compared to the static 2D model. This could be because the 2D model’s rigid base substrate and static culture technique put the cells under stress, mimicking signs of endothelial dysfunction. Given that these results corroborate the theory that relative arginase hyperactivity causes endothelial dysfunction in mice (Vaisman et al., 2012) and in diabetes patients with ED condition (Kövamees et al., 2016). Furthermore, the loss of fluid flow combined with a mTOR-based regulatory mechanism to balance eNOS and arginase II expression results in elevated ^13^C_5_, ^15^N_2_ L-Ornithine and decreased ^13^C_6_ ^15^N_3_ L-citrulline levels, which indicates ED, was reported in these studies (Decker and Pumiglia, 2018; Mammedova et al., 2021).

Flow induced citrulline and NO production was significantly higher compared to static culture was reported by Noris et.al group (Noris et al., 1995). Our findings also show higher citrulline levels under flow conditions with the reason that fluid shear stress induce the signal of PECAM-1 that act through adaptor molecules such as phosphatidylinositol-3-kinase (PI3K), which then activate eNOS, mTOR and the transcription factor Klf2 to regulate the functional genes (Zhou et al., 2014). Therefore, lower ratio in 2D static culture mimics more of endothelial dysfunction conditions.

To understand the impact of fluid flow on the substrate availability, we calculated the arginine bioavailability using the formula ^13^C_6_,^15^N_4_ L-arginine /(^13^C_6_ ^15^N_3_ L-citrulline + ^13^C_5_, ^15^N_2_ L-Ornithine) **(Fig. 5G)**. The result obtained signify that the 2D platform delivers lower arginine bioavailability compared to the 3D flow models. Such measurements aid in the comprehension of arginine intake and the efficient conversion that occurs in 2D as compared to 3D. Nonetheless, the 3D model’s crucial readout that indicates the healthy status with the balanced ratio. In summary, the analysis demonstrates that the lower ^13^C_6_ ^15^N_3_ L-citrulline /^13^C_5_, ^15^N_2_ L-Ornithine ratio, and reduced L-arginine bioavailability in static culture may be related to underlying vascular dysfunction due to enhanced “catabolic pathway” of L-arginine, in particular more of L-ornithine production, resulting in less L-arginine consumption as a substrate for NO generation.

## 4. Conclusion

Our research sheds light on the influence of fluid shear stress on endothelial function at a metabolic level, employing our innovative 3D microvessel-on-a-chip model with a unidirectional flow system generated by a microfluidic pump. Additionally, our optimized MS method proves to be ideal for measuring NO marker metabolites at both extracellular and intracellular levels, providing valuable insights into metabolic flux. In future studies, this method could be applied to assess the levels of NO marker metabolites in patient profiles by perfusing patient plasma samples in our 3D model and analyzing the degree of pathological conditions and drug responses. Combining flux data with other omics data in future studies would offer a more comprehensive understanding of NO’s vasodilatory effects in studying endothelial dysfunctions. Furthermore, enhancing the pump system with higher shear forces could be explored to apply to various endothelial cells types, emulating different microvessel models. In conclusion, our validation of shear stress’s impact on eNOS phosphorylation using our 3D cell culture model from a metabolic perspective is a novel approach that confirms the physiological status.

## Author Contributions

Kanchana Pandian, Conceptualization, Investigation, Methodology, Writing - original draft; Luojiao Huang, Methodology – mass spectrometry method, Writing – review and editing; Abidemi Junaid, Methodology – microfluidic pump designing, Writing – review and editing; Amy Harms, Writing – review and editing;

Anton Jan van Zonneveld and Thomas Hankemeier, Conceptualization, Resources, Supervision, Funding acquisition, Writing – review and editing.

## Acknowledgements

This project has received funding from the European Union’s Horizon 2020 research and innovation program (LogicLab network) under the Marie Skłodowska-Curie grant agreement No 813920. Part of these studies we supported by the Dutch Heart Foundation (CVON RECONNECT) and ZonMW (MKMD: 114022501) grants to T.H and A.J.V.Z.

