## supplementary file for "Tracer-based metabolomics for profiling nitric oxide metabolites in a 3D microvessel-on-a-chip model"

**Figure 1**

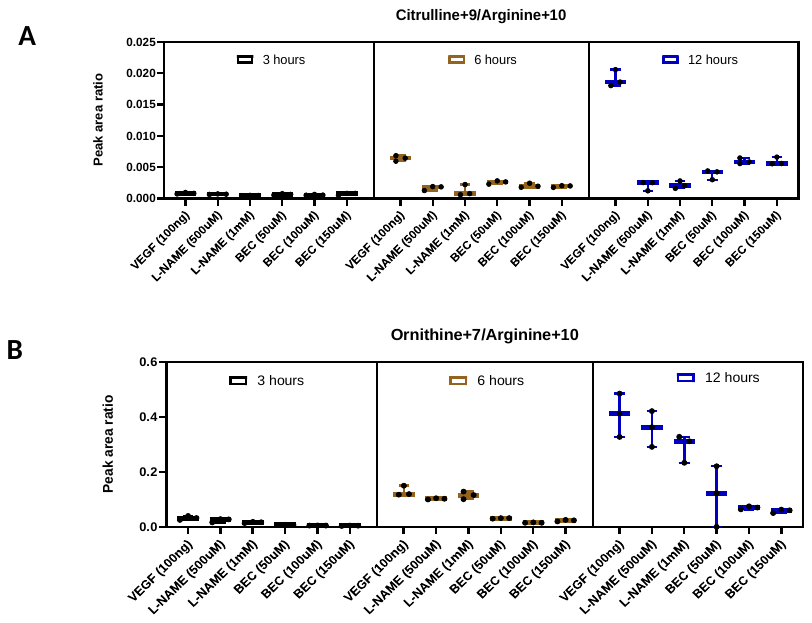

**Supplementary Fig 1. Dosage study of stimulatory and inhibitory compounds and the extracellular measurement of marker metabolites.**

(A & B) Ratios of ^13^C_6_, ^15^N_3_ L-Citrulline (Citrulline+9), ^13^C_5_, ^15^N_2_ L-Ornithine (L-Ornithine+7) and ^13^C_6_, ^15^N_4_-L-arginine (L-Arginine+10) after the different dosage treatment of stimulator and inhibitors of eNOS and arginase enzyme on HCAECs at different incubation hours. Incubation hours represented in different colors. 3 hours – black bars; 6 – hours – brown bars; and 12 hours – blue bars. Significance determined by one way ANOVA multiple comparison test. For Citrulline+9/Arginine+10 and for Ornithine+7/Arginine+10, overall significant difference was observed between 3, 6 and 12 hours with *p* value <0.0001.

**Figure 2**

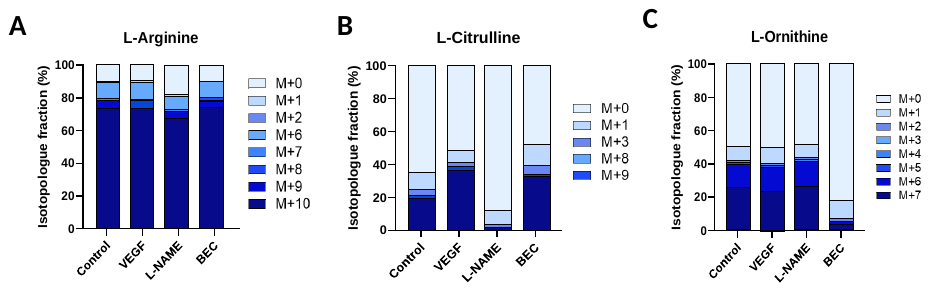

**Supplementary Fig 2.** **Isotopologue fractions** of (A) L-Arginine (B) L-Citrulline and (C) L-Ornithine in stimulated and inhibited treatments and in control.

**Figure 3**

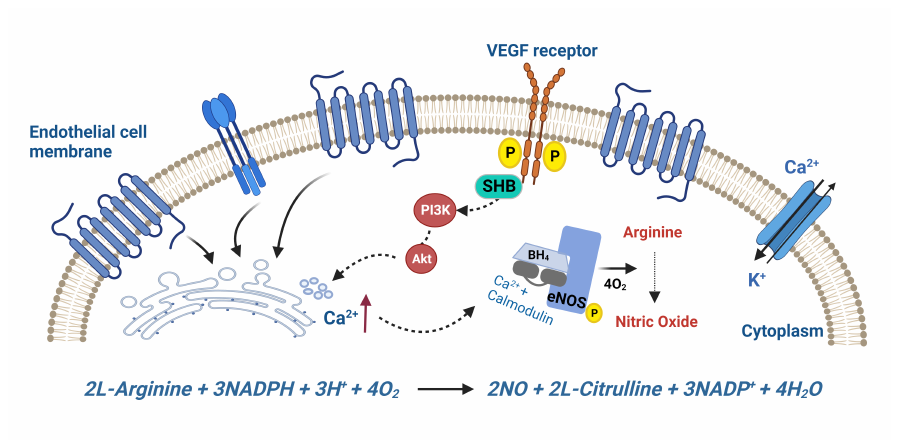

**Supplementary Fig. 3. Schematic representation of Vascular Endothelial Growth Factor (VEGF) stimulation over Enos enzyme and NO production.**

The external signals – agonist of nitric Oxide (NO) such as acetylcholine, ATP & ADP (adenosine triphosphate and diphosphate),VEGF etc., has specific receptor cite on the endothelial cell surface (Blue). When VEGF binds to the muscarinic receptor, it releases Calcium2+, a major factor that leads to the chain of events cause NO production. Once the receptors are stimulated by the agonists (VEGF) they act upon the endoplasmic reticulum (ER) by phosphorylating the VEGF receptor and transfer the signal via SHB adaptor protein, PI3K (Phosphoiniositide3 kinase) and protein kinases (Akt), increase Ca2+ level, that binds to the calmodulin protein and cause some conformational changes. In the presence of tetrahydrobiopterin (BH4) and oxygen it phosphorylate the enhance the Enos phosphorylation. The activated Enos will convert L-Arginine to L-Citrulline and Nitric Oxide (NO). Calcium level is balanced by the calcium potassium channel from the circulation. Illustration created BioRender.com, accessed on HV24G0AUF2, September 24, 2022.

**Figure 4**

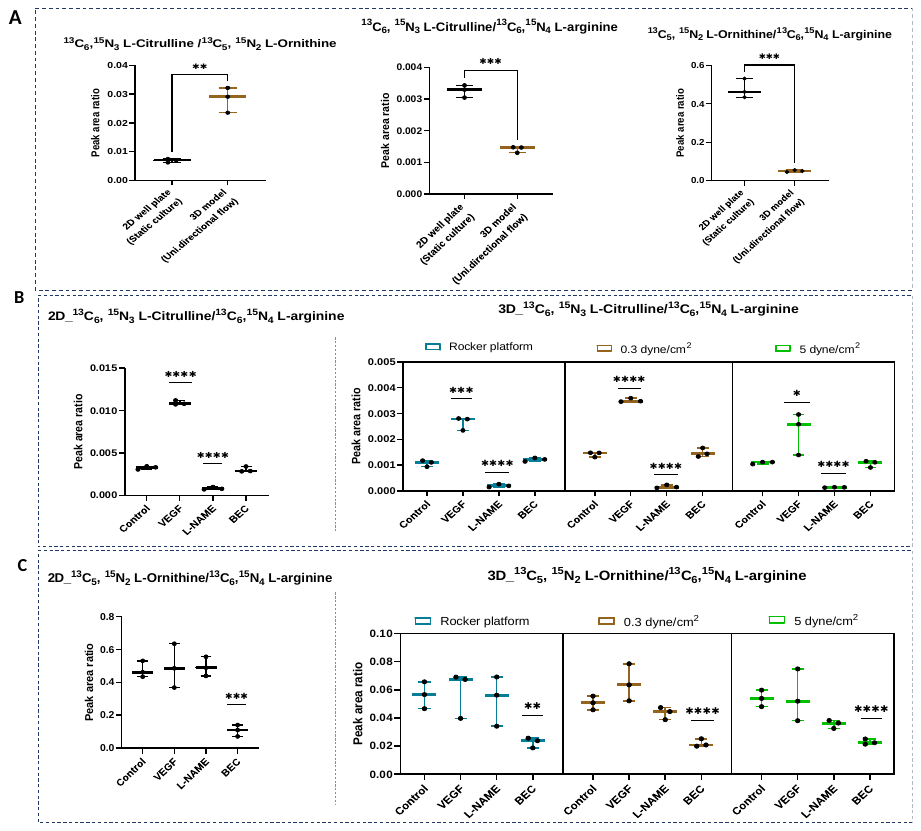

**Supplementary fig. 4. Measurement of extracellular marker metabolites in all platforms.** A) Comparison of metabolites ratio between 2D and 3D (0.3 dyne/ cm^2^) model platforms in controls B) Ratio of ^13^C_6_,^15^N_3_ L-Citrulline / ^13^C_6_,^15^N_4_ L-arginine C) Ratio of ^13^C_5_,^15^N_2_ L-Ornithine / ^13^C_6_,^15^N_4_ L-arginine in stimulatory and inhibitory compounds treated samples from 2D static, 3D rocker platform (bidirectional flow model) & the rerouted OrganoPlate with microfluidic perfusion pump operated with different shear stresses (unidirectional flow model).

Metabolic data significance determined by one way ANOVA multiple comparison test. *, P<0.05; **, P<0.01; ***, P <0.001; ****, P<0,0001*.*

**Supplementary table. 1.** Modified MS acquisition method of 29 transition pairs of potential isotopologues of marker metabolites.

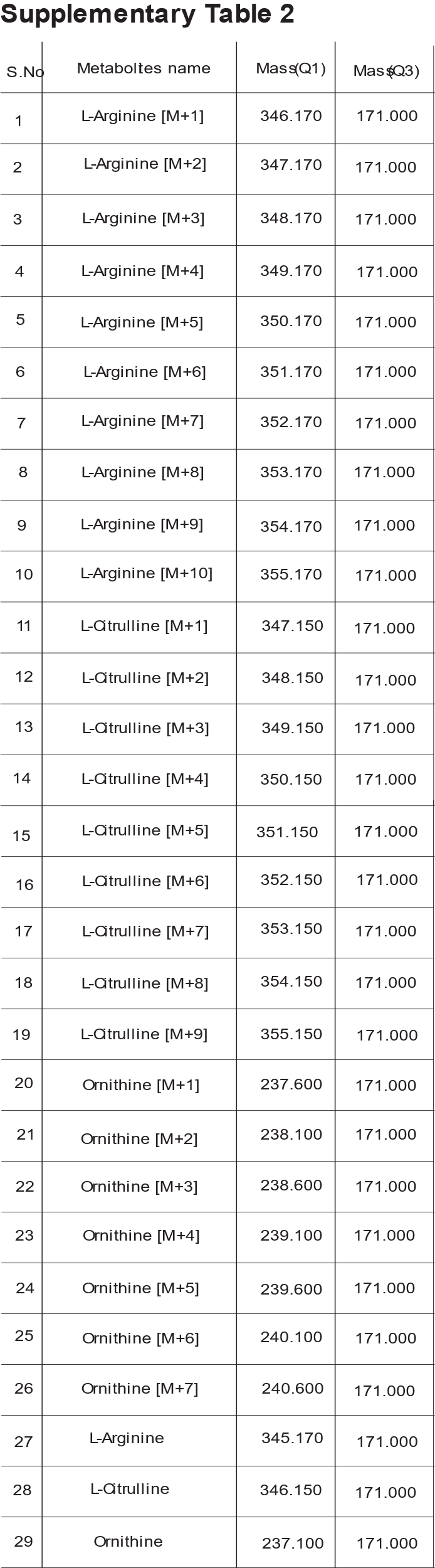
